# YAP/TAZ signaling in allantois-derived cells is required for placental vascularization

**DOI:** 10.1101/2024.09.15.613151

**Authors:** Siqi Gao, Triloshan Thillaikumaran, Martin H. Dominguez, William Giang, Kevin Hayes, Xiaowen Chen, Jesse Pace, Jenna Bockman, Danielle Jathan, Derek Sung, Sweta Narayan, Maxwell Frankfurter, Patricia Mericko-Ishizuka, Jisheng Yang, Marco Castro, Michael Potente, Mark L. Kahn

**Affiliations:** Cardiovascular Institute, Department of Medicine, Perelman School of Medicine, University of Pennsylvania, Philadelphia, Pennsylvania, USA; Advanced Light Microscopy Core, Pennsylvania State University College of Medicine, Hershey, Pennsylvania, USA; Berlin Institute of Health (BIH) and Charité-Universitätsmedizin Berlin, Berlin, Germany

**Keywords:** Hippo signaling, placenta, endothelial cells, stromal cells, lightsheet

## Abstract

Normal placental development and angiogenesis are crucial for fetal growth and maternal health during pregnancy. However, molecular regulation of placental angiogenesis has been difficult to study due to a lack of specific genetic tools that isolate the placenta from the embryo and yolk sac. To address this gap in knowledge we recently developed *Hoxa13*^*Cre*^ mice in which Cre is expressed in allantois-derived cells, including placental endothelial and stromal cells. Mice lacking the transcriptional regulators Yes-associated protein (YAP) and PDZ-binding motif (TAZ) in allantois-derived cells exhibit embryonic lethality at midgestation with compromised placental vasculature. snRNA-seq analysis revealed transcriptional changes in placental stromal cells and endothelial cells. YAP/TAZ mutants exhibited significantly reduced placental stromal cells prior to the endothelial architectural change, highlighting the role of these cells in placental vascular growth. These results reveal a central role for YAP/TAZ signaling during placental vascular growth and implicate *Hoxa13*-derived placental stromal cells as a critical component of placental vascularization.

## Introduction

The vascular system functions as a critical component for organ development^1,2^. The mammalian placenta is a highly vascularized organ, which enables exchange of nutrients, gases, and wastes between mother and fetus. The allantois, a site distinct from the embryo and yolk sac, is responsible for the development of the fetal placental vasculature^3^. However, the signaling pathways that regulate placental growth remain largely unknown.

Vascular development is tightly regulated by transcriptional programing^4^. YAP/TAZ are transcriptional regulators in the Hippo signaling pathway that modulate gene expression in response to diverse extracellular stimuli, including mechanical force, cell-cell contact, and soluble ligands^5-7^. Several studies have found that YAP/TAZ signaling plays versatile roles in different vascular beds^8-13^. The role of YAP/TAZ in the placental vasculature has not been explored previously.

In the present study, we utilize the Cre-expressing mouse line *Hoxa13*^*Cre*^, which targets allantois-derived cells, to demonstrate that the transcriptional regulators YAP/TAZ in these cells are indispensable for the developing placenta and embryonic survival. Specifically, YAP/TAZ are required for the expansion of placental stromal cells in the *Hoxa13*^*Cre*^ lineage and for placental vascularization. Together, these findings highlight a previously unreported role for YAP/TAZ in placental vascularization and development.

## Results

### Deletion of YAP/TAZ in allantois-derived cells impairs fetal vascular growth in the placenta and results in embryonic lethality

To investigate the role of YAP and TAZ in the developing placenta, we first assessed expression of their mRNA transcripts and proteins in the placental labyrinth. Both YAP and TAZ are widely expressed across a range cell types within this region (Supplemental Figure1), including vascular endothelial cells and stromal cells. This broad expression pattern suggests that YAP/TAZ may play a significant role during the development of the placenta.

To test the role of YAP/TAZ signaling during placental vascular development, we crossed *YAP*^*fl/fl*^*;TAZ*^*fl/fl*^ animals to the *Hoxa13*^*Cre/+*^ allele. The *Hoxa13*^*Cre/+*^ allele is active in the early allantois and lineage tracing studies reveal recombination in virtually all placental endothelial and stromal cells ^14^ (Supplemental Figure2). Intercrosses of *YAP*^*fl/fl*^*;TAZ*^*fl/fl*^ and *YAP*^*fl/+*^*;TAZ*^*fl/+*^; *Hoxa13*^*Cre/+*^ mice revealed no *YAP/TAZ*^*fl/fl*^; *Hoxa13*^*Cre/+*^ animals at weaning (Table 1, expected at 12.5%, P<0.0021). These results indicate that expression of YAP/TAZ in allantois-derived cells is crucial for embryonic survival.

**Table 1:** Live offspring at weaning from *YAP*^*fl/fl*^*;TAZ*^*fl/fl*^ X *YAP*^*fl/+*^ *;TAZ*^*fl/+*^; *Hoxa13*^*Cre*^ crosses. *YAP*^*fl/fl*^*;TAZ*^*fl/fl*^ females were crossed with *YAP*^*fl/+*^*;TAZ*^*fl/+*^; *Hoxa13*^*Cre*^ males, and live progeny from litters were genotypes at weaning. No *YAP*^*fl/fl*^*;TAZ*^*fl/fl*^; *Hoxa13*^*Cre*^ mice were recovered [χ^2^ (7dof)=24.41, p<0.002].

| Genotype | Observed # (%) | Expected # (%) |
| --- | --- | --- |
| $YAP^{fl/+}; TAZ^{fl/+}$ | 17 (21.8) | 9.75 (12.5) |
| $YAP^{fl/fl}; TAZ^{fl/+}$ | 9 (11.5) | 9.75 (12.5) |
| $YAP^{fl/+}; TAZ^{fl/fl}$ | 15 (19.2) | 9.75 (12.5) |
| $YAP^{fl/fl}; TAZ^{fl/fl}$ | 14 (17.9) | 9.75 (12.5) |
| $YAP^{fl/+}; TAZ^{fl/+}; Hoxa13^{Cre/+}$ | 8 (10.3) | 9.75 (12.5) |
| $YAP^{fl/fl}; TAZ^{fl/+}; Hoxa13^{Cre/+}$ | 0 (0) | 9.75 (12.5) |
| $YAP^{fl/+}; TAZ^{fl/fl}; Hoxa13^{Cre/+}$ | 15 (19.2) | 9.75 (12.5) |
| $YAP^{fl/fl}; TAZ^{fl/fl}; Hoxa13^{Cre/+}$ | 0 (0) | 9.75 (12.5) |

We next performed timed-matings to assess when *YAP/TAZ*^*fl/fl*^; *Hoxa13*^*Cre/+*^ embryos die. At E10.5, both *YAP/TAZ*^*fl/fl*^ and *YAP/TAZ*^*fl/fl*^; *Hoxa13*^*Cre/+*^ embryos were viable and appeared comparable. In contrast, at E11.5 YAP*/TAZ*^*fl/fl*^; *Hoxa13*^*Cre/+*^ mutant embryos were pale and exhibited a slow heart rate (Figure 1). All *YAP/TAZ*^*fl/fl*^; *Hoxa13*^*Cre/+*^ embryos were dead at E12.5 and resorbed by E13.5. *YAP/TAZ*^*fl/fl*^; *Hoxa13*^*Cre/+*^ placentas were significantly smaller compared to littermate controls at E13.5 (Figure 1). To evaluate the placental vascular labyrinth, we performed H&E staining of tissue placenta sections from E10.5 to E13.5 in littermates. At E10.5, *YAP/TAZ*^*fl/fl*^ and *YAP/TAZ*^*fl/fl*^; *Hoxa13*^*Cre/+*^ placentas exhibited a similar labyrinth area. From E11.5, labyrinth area is significantly reduced in *YAP/TAZ*^*fl/fl*^; *Hoxa13*^*Cre/+*^ placentas (Figure 1). Notably, the fetal labyrinth vessels are significantly dilated in *YAP/TAZ*^*fl/fl*^; *Hoxa13*^*Cre/+*^ compared with controls at this time-point (Figure 1).

**Figure 1.**
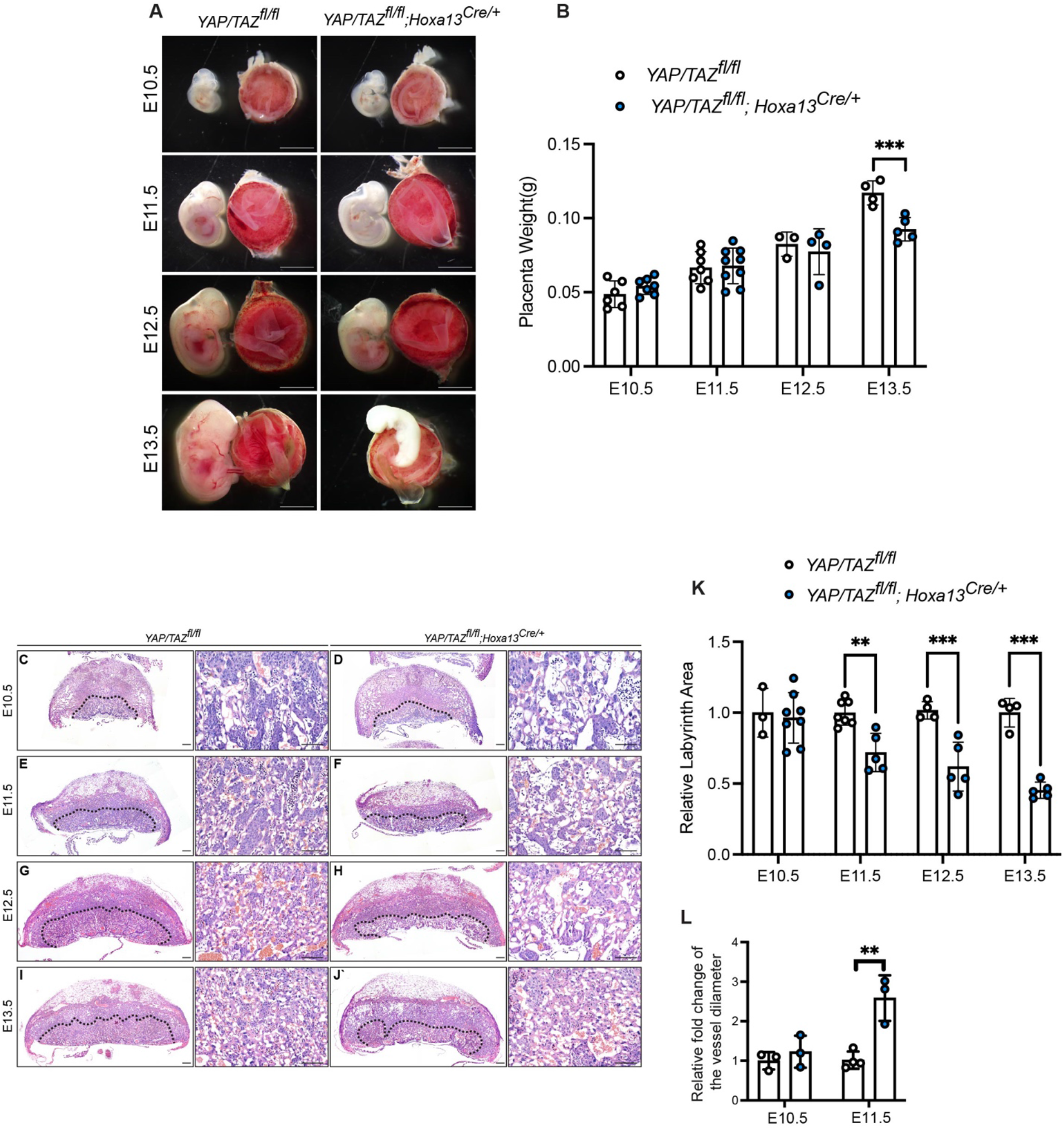
YAP/TAZ deletion in allantois derived cells results in embryonic lethality and reduced placental labyrinth area. (A) Representative gross morphology images from E10.5 to E13.5 Control (*YAP/TAZ*^*fl/fl*^) and *YAP/TAZ*^*fl/fl*^; *Hoxa13*^*Cre/+*^. (B) Placentae were weighed from timed-mating dissections. Data are expressed as mean + SEM. *** p < 0.001. n = 4-9 mice per group. Statistical analyses were performed using two-tailed Student’s *t*-test. Scale bars: 1mm. (C-J) Hematoxylin and eosin (H&E) staining from E10.5 to E13.5 Control (*YAP/TAZ*^*fl/fl*^) and *YAP/TAZ*^*fl/fl*^; *Hoxa13*^*Cre/+*^ serial placenta sections. Dotted lines denoted for the labyrinth area. Scale bars: 50 μm. (K) Quantification of the relative labyrinth area. (L) Quantification of labyrinth vessel diameter. Each dot represents a single animal.

To better assess the impact of YAP/TAZ deletion on placental labyrinth vascular morphology, we adapted the iDISCO protocol to clear the placenta and image it using light-sheet microscopy^15^. Labyrinth vascular volume and structure were comparable in *YAP/TAZ*^*fl/fl*^ and *YAP/TAZ*^*fl/fl*^; *Hoxa13*^*Cre/+*^ placentas at E10.5. However, at E11.5 labyrinth vascular volume was significantly reduced in *YAP/TAZ*^*fl/fl*^; *Hoxa13*^*Cre/+*^ placentas compared with control (Figure 2). In addition, the placental vascular network appeared highly dysmorphic in *YAP/TAZ*^*fl/fl*^; *Hoxa13*^*Cre/+*^ mutants, with regions lacking vasculature as well as smaller caliber vessels (N= 3-4 for each genotype, Figure 2).

**Figure 2.**
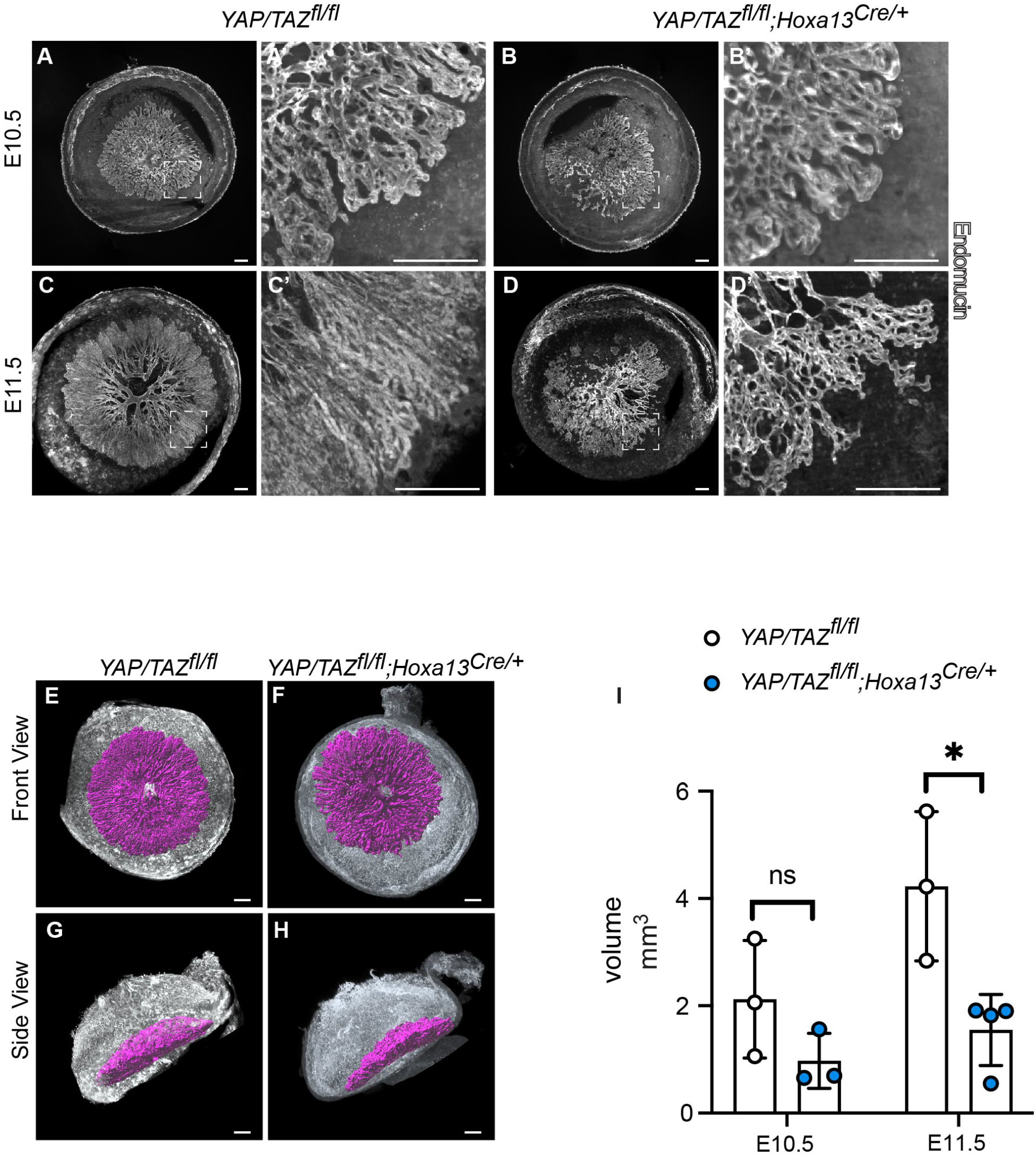
Light-Sheet Fluorescence Microscopy analysis of *YAP/TAZ*^*fl/fl*^; *Hoxa13*^*Cre/+*^ placental vasculature. (A-D) E10.5 and E11.5 control and *YAP*^*fl/fl*^*;TAZ*^*fl/fl*^; *Hoxa13*^*Cre/+*^ placenta tissues were whole mount stained with endomucin and subjected to clearing and then imaged by UltraMicroscope II. (A’-D’) inserts of (A-D) the labyrinth region. Scale bars: 300μm. (E-H) 3D rendering of segregated E11.5 Labyrinth area. Scale bars: 500μm. (I) Quantification of vascular volume is generated via Imaris software. *p < 0.05 as calculated by Student’s *t* test.

In addition, histological analysis of E11.5 *YAP/TAZ*^*fl/fl*^; *Hoxa13*^*Cre/+*^ embryos revealed hearts with a thinned myocardium (Supplemental Figure 3). Since Hoxa13Cre is not active in the embryonic heart^14^, the heart defects in *YAP/TAZ*^*fl/fl*^; *Hoxa13*^*Cre/+*^ are secondary to placental defects. This observation is consistent with emerging evidence that defects in the placenta can lead to embryonic lethality and the etiology of congenital heart defects^16,17^.

### YAP/TAZ are required for placental stromal cells to facilitate vascularization

To gain a deeper understanding of the cellular and molecular changes in *YAP/TAZ*^*fl/fl*^; *Hoxa13*^*Cre/+*^, we isolated single nuclei from control and *YAP/TAZ*^*fl/fl*^; *Hoxa13*^*Cre/+*^ mutants and carried out single-nuclei RNA sequencing (snRNA-seq) using the 10x Genomics platform. An integrated analysis revealed distinct clusters of endothelial cells and mesenchymal stromal cells (Figure 3, Supplemental Figure 4). All samples were identified to contain the same cell populations (Supplemental Figure 4). In the mesenchymal stromal cell clusters, *Acta2*^*+*^ populations are reduced in *YAP/TAZ*^*fl/fl*^; *Hoxa13*^*Cre/+*^ mutants (Figure 3). Comparing mutant and control populations of fetal mesenchymal stromal cells, mutant stromal cells revealed a significant increase in expression of 58 genes, and significant reduction in the expression of 33 genes. Gene ontology (GO) analysis of biological processes in mesenchymal stromal cells revealed that labyrinthine layer morphogenesis, positive regulation of smooth muscle cell proliferation, cell adhesion and extracellular organization were enriched. The endothelial cell population exhibits 20 significantly increased genes and 19 decreased genes. Regulation of smooth muscle contraction and migration, blood vessel remodeling, and cell adhesion are enriched in GO analysis of biological processes in ECs (Figure 3).

**Figure 3.**
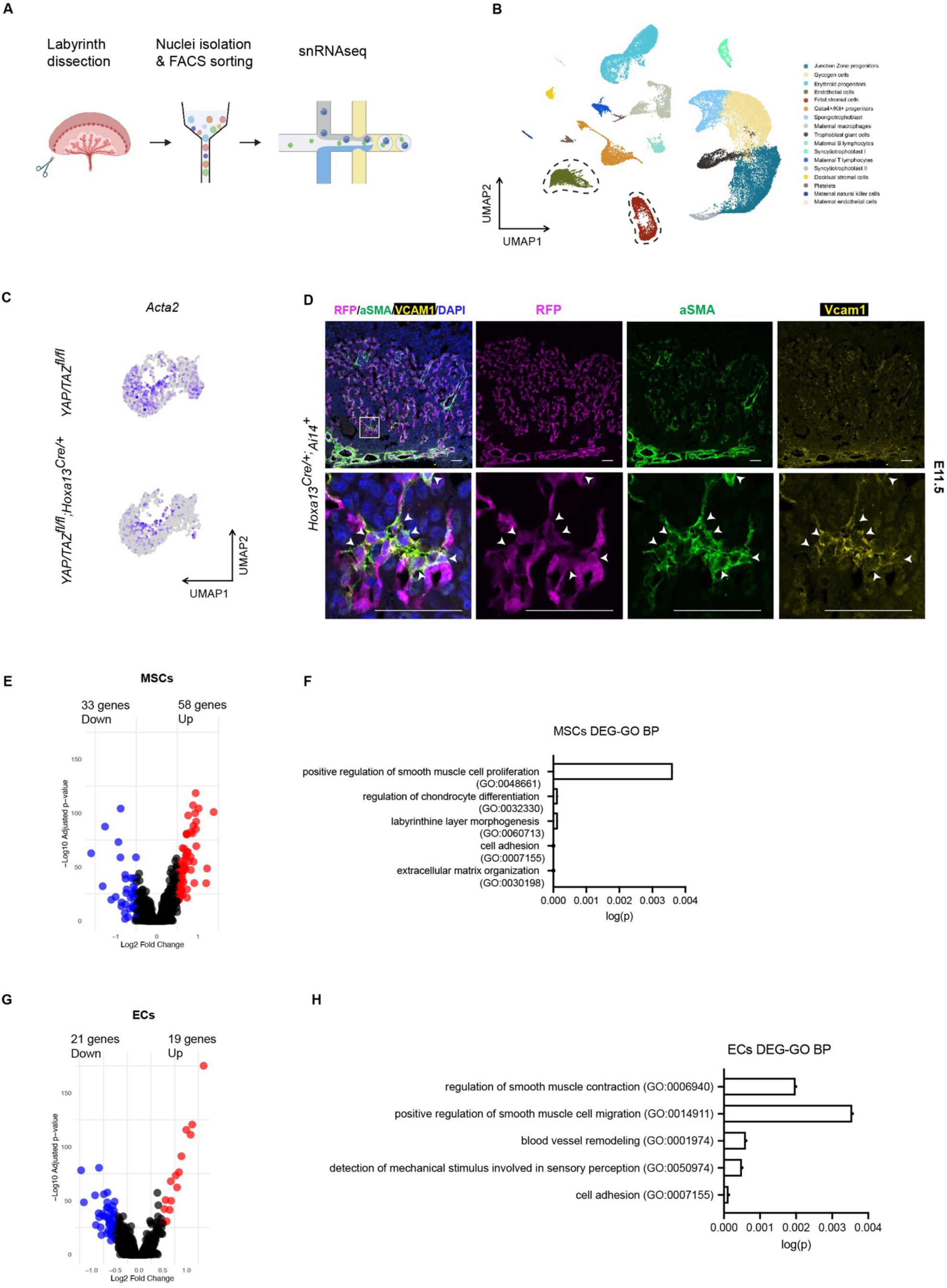
snRNAseq analysis reveal transcriptional changes in *YAP*^*fl/fl*^*;TAZ*^*fl/fl*^; *Hoxa13*^*Cre/+*^ placentas. (A) Schematic workflow of generating placental snRNA-seq dataset from control (*YAP*^*fl/fl*^*;TAZ*^*fl/fl*^) and *YAP*^*fl/fl*^*;TAZ*^*fl/fl*^; *Hoxa13*^*Cre/+*^ E10.5 and E11.5 placentas. (B) UMAP dimension reduction analysis of snRNAseq data shows distinct cell clusters in the placental labyrinth. (C) *Acta2*^*+*^ (aka. αSMA) stromal cells in control and *YAP*^*fl/fl*^*;TAZ*^*fl/fl*^; *Hoxa13*^*Cre/+*^ placentas. (D) Representative images of *Hoxa13*^*Cre*^ labels VCAM1^+^/αSMA^+^ stromal cells in the E11.5 placenta. Immunofluorescence staining for RFP (magenta), αSMA(green), VCAM1 (yellow), and DAPI (blue). Scale bars: 50 μm. (E and G) Volcano plots show relative expression of mesenchymal stromal cells (MSCs) and endothelial cells (ECs); Red points represent significantly upregulated genes (log2FC > 0.5 and padj < 0.05), blue points represent significantly downregulated genes (log2FC < -0.5 and padj < 0.05), and black points are genes that are not significantly differentially expressed. Fetal mesenchymal stromal cell cluster is defined by robust expression of *Col1a2* and *Angpt1*. Endothelial cell cluster is defined by *Kdr* and *Tek* (Supplemental Figure 4). (F and H) Gene ontology (GO) biological process (GO) enrichment performed for mesenchymal stromal cells (F) and endothelial cells (H).

Next, to determine whether placental *Acta2*^*+*^ mesenchymal stromal cells are affected in the *YAP/TAZ*^*fl/fl*^; *Hoxa13*^*Cre/+*^ placenta, we directly examined α-smooth muscle actin (αSMA) expression using immunostaining of placental tissue sections. Analysis of *Hoxa13-Cre;Ai14* placentas in which Cre activity is marked by expression of the TdTomato red fluorescent reporter revealed that *Acta2*^*+*^ mesenchymal stromal cells are also derived from *Hoxa13*^*+*^ lineage, and co-express VCAM1 (Figure 3). Placental VCAM1^+^ stromal cells have been shown to possess pro-angiogenic activity^18^. *YAP/TAZ*^*fl/fl*^; *Hoxa13*^*Cre/+*^ placentas exhibited significantly reduced αSMA^+^/VCAM1^+^ stromal cells surrounding the fetal blood vessels at the labyrinth at E10.5, a timepoint prior to the most significant observed changes in vascular volume (Figure 4). At E11.5, significant low coverage of αSMA^+^/VCAM1^+^ stromal cells was observed in *YAP/TAZ*^*fl/fl*^; *Hoxa13*^*Cre/+*^ placental labyrinth and an absence of stromal cells at the chorioallantoic plate was observed (Figure 4 and Supplemental Figure 5). These data suggest that YAP/TAZ signaling may be required for the expansion of *Hoxa13Cre* lineage stromal cells during placental vascularization.

**Figure 4.**
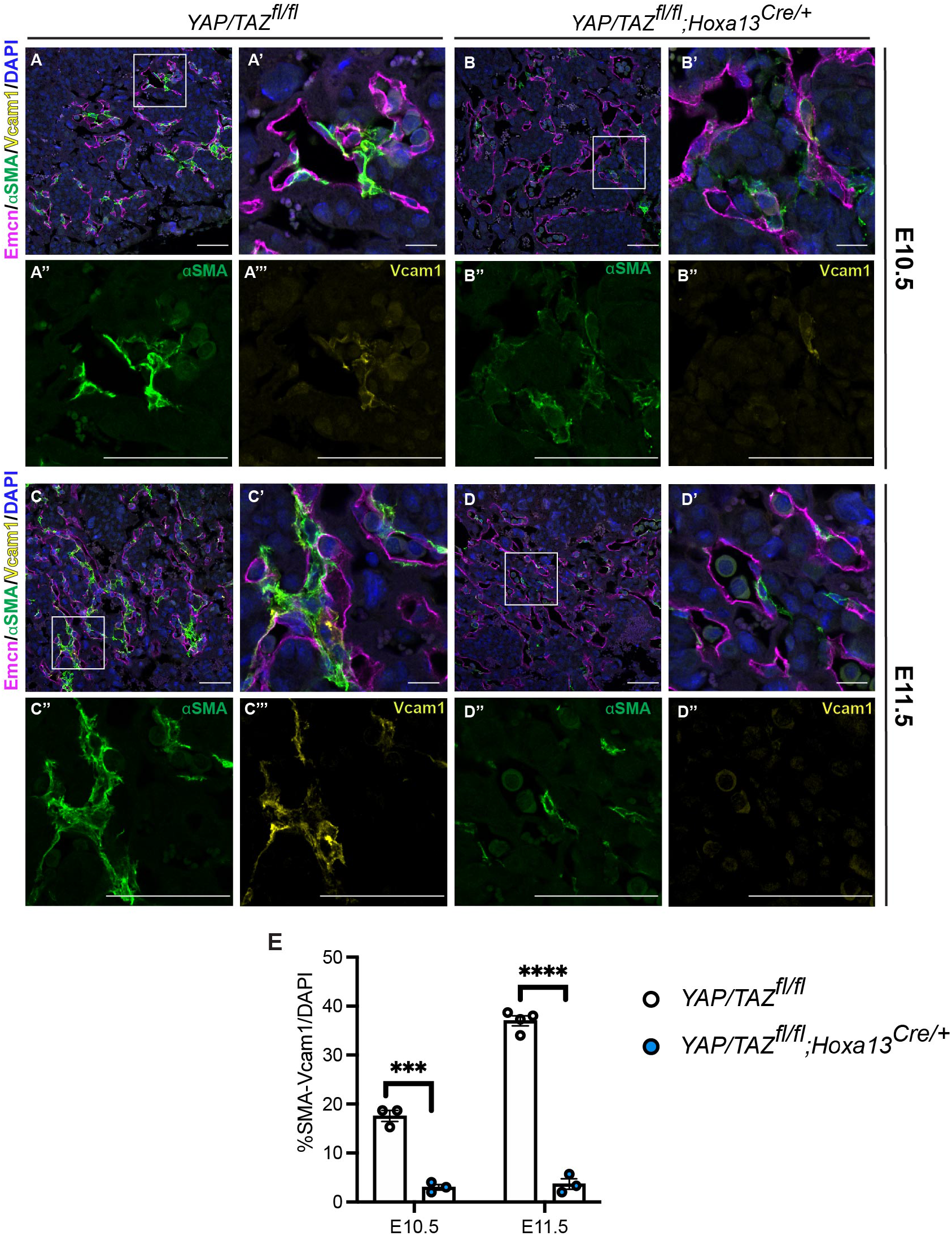
*YAP/TAZ*^*fl/fl*^; *Hoxa13*^*Cre/+*^ exhibits reduced stromal cells in the placenta labyrinth. (A-D) Sections from E10.5 to E11.5 control and *YAP/TAZ*^*fl/fl*^; *Hoxa13*^*Cre/+*^ were immunostained for endothelial cell marker endomucin (magenta), stromal cell marker αSMA (green) and Vcam1 (yellow), and nuclei were counterstained with Hoechst (blue). (A’-D’) inserts of (A-D) the labyrinth region. Scale bars: 50μm. (E) Quantification of the percentage αSMA/VCAM1 positive cells present at the labyrinth area. ***p < 0.001 as calculated by Student’s *t* test.

## Discussion

Allantois-derived cells are known to form the fetal placental vasculature^3^, yet the molecular players underlying this placental vascular growth remain unclear. We demonstrate that loss of YAP/TAZ in *Hoxa13*^*+*^ lineage cells leads to profound defects in placental vascularization and in the expansion of placental stromal and endothelial cells.

Our studies implicate a role for YAP/TAZ signaling in placental endothelial cells and/or stromal cells for successful vascular growth. YAP/TAZ signaling has been shown to drive angiogenesis in the retina through activation of nutrient-driven mTORC signaling^10^, while in vitro studies have connected the pathway to endothelial proliferation and common angiogenic signaling mechanisms such as VEGF and angiopoietin signaling^8,13^. In addition, prior studies have shown that YAP/TAZ signaling contributes to the emergence of neural crest-derived vascular smooth cells at the branchial arches^19,20^, and contractile phenotype of vascular smooth muscle cells^21^. Our studies identify defects in both endothelial cells and stromal cells, but we observed that stromal cell abnormalities in *YAP/TAZ*^*fl/fl*^; *Hoxa13*^*Cre/+*^ mutants appear prior to significant differences in vascular area and structure (Figure 4). Thus, the primary defect may be loss of stromal cell signaling required to support placental vascular growth. Similar with loss of either PDGFB or PDGFRβ leads to dilated fetal blood vessels^22^, here we found that αSMA^+^/VCAM1^+^ stromal cells affect placental labyrinth fetal vessel diameters and vascularization.

Our study also highlights remaining limitations for investigation of placental vascular growth. Constitutive deletion driven by the Hoxa13^Cre^ line in the early allantois confers genetic deletion in multiple cell types in the placenta, including both placental stromal and endothelial cells. Future studies will require second generation tools, e.g. a Dre-Cre intersectional genetic strategy, to limit genetic manipulation to a single cell type as well as tools to temporally control genetic changes.

## Acknowledgement

We thank Kahn lab members for helpful discussions and technical advice and the CDB Microscopy Core for support with microscopy. We thank the Advanced Light Microscopy Core at Penn State College of Medicine for support with light sheet imaging.

## Author Contributions

SG designed and performed most of the mouse experiments. TT, XC, MD, KH, JB, DJ, SN, JP, DS, MF, WG and PM contributed to mouse studies. JY performed histological studies. SG and MLK interpreted data and wrote the manuscript.

## Data and Materials Availability

All data and reagents will be made available upon reasonable request. Transgenic mouse lines are available from Dr. Mark Kahn under a material transfer agreement with the University of Pennsylvania.

## Materials and Methods

### Animals

YAP/TAZ-floxed mice (YAP/TAZ^fl/fl^), and Hoxa13-Cre transgenic mice have been described^23^ (JAX cat# #030532). All mice were maintained on a mixed genetic background at the University of Pennsylvania animal facility. YAP/TAZ-floxed and Hoxa13^Cre^ transgenic embryos and mice were genotyped as described^14,23^. All procedures were conducted using an approved animal protocol (806811) in accordance with the University of Pennsylvania Institutional Animal Care and Use Committee.

### Histological and Immunofluorescence Staining

For paraffin slides, tissue sections were deparaffinized and rehydrated through xylene and ethanol gradients. Sections were then subjected to Hematoxylin and Eosin (H&E) staining or processed for immunofluorescence. H&E staining was performed on placental and embryonic sections as previously described^24^. For immunofluorescence, pH6.0 citrate antigen retrieval buffer (Sigma-Aldrich, C9999) were used. Sections were incubated in the primary antibody diluted in diluent buffer (IHC World, Cat# IW-1000) overnight at 4°C. Sections were then washed three times with PBS, then incubated at room temperature for 1 hour in secondary antibody at a 1:500 concentration in PBS. For cryosectioned slides, sections were taken from the freezer before staining and warmed at room temperature for 2 minutes, and washed once with PBS, then incubated in the primary and secondary antibody as described above.

Primary antibodies used for immunofluorescence: Endomucin (1:300, R&D#AF4666), aSMA-Cy3 (1:300, Sigma#C6198), Vcam1 (1:200, R&D#AF643), RFP (1:200, Rockland#600-401-379), YAP (1:100, CellSignaling#14074S), TAZ(1:100,Sigma#HPA007415), MCT1 (1:200, Millipore#AB1286-I), MCT4 (1:200,Millipore#AB3314P).

Images were obtained with a Zeiss LSM 710 confocal microscope,Zeiss Axio Observer 7 widefield microscope, and an Olympus BX53 widefield fluorescence microscope. Images were processed, visualized, and analyzed using ImageJ/FIJI software^25^.

### Whole mount immunolabeling and clearing

Tissue samples were fixed with 4% PFA overnight. And samples were washed with PBS for twice (15 minutes each) and permeabilized with 0.5% Triton X-100 at room temperature overnight. Samples were incubated in the 1% normal donkey serum blocking buffer at room temperature overnight, followed by incubating in the primary antibody (Endomucin, 1:300, R&D AF4666) at room temperature for 3 days. Samples were washed with PBST four times (30 minutes each). Samples were next incubated in the diluted secondary antibody at 4C for 2 days. Tissue samples were then washed with PBST five times (30 minutes each). Stained tissues were subjected to clearing with modified iDISCO protocol^15,26^. Briefly, tissue samples were dehydrated with gradient methanol series (20%, 40%, 60%, 80%, 100%; 1 hour each). Samples were incubated in 66% DCM (Dichloromethane, Sigma 270997) / 33% methanol at room temperature for 3 hours with shaking following by incubating with 100% DCM twice (15 minutes each). Tissue samples then were incubated in 100% Dibenzyl ether until tissue is clear.

### Light-sheet imaging

Tissue samples were imaged with a Miltenyi BioTec Ultramicroscope II light-sheet microscope in a quartz cuvette filled with dibenzyl ether. Images were acquired with ImSpector Pro 7.1.16 software and had an isotropic voxel size (2.71 μm x 2.71 μm x 2.71 μm. We used a 4x/0.35 NA MI PLAN objective with a dipping cap for organic solvents. All six sheets were used with the following light-sheet parameters: 100% width and 0.026 sheet NA. Volume quantification of light sheet images was performed through Imaris v10.0.

### Single nuclei RNA-sequencing

Nuclei were isolated from snap frozen labyrinth tissue samples following the 10X nuclei isolation kit (10X# 1000494). Isolated nuclei were labeled with DAPI and RNase inhibitor, and were purified using FACS (MoFlo Astrios Sorter, 100μm nozzle) and collected in isolation buffer (containing RNase inhibitor 0.2 U/μl). Sorted nuclei were subsequently loaded onto the 10X Genomics Chromium Controller. mRNA will be collected and converted to snRNA-seq libraries using 10X Single Cell 3’ v3 kit, and high-throughput sequencing will be performed on an Illumina NovaSeq6000. The gene expression matrices for each dataset were generated using the CellRanger cloud analysis. The counts matrix was thresholded and analyzed in the package Seurat (v 4.3.0). Cells were first filtered to have >500 detected genes and less than 15% of total UMIs mapping to the mitochondrial genome. Dimension reduction was performed using the uniform manifold approximation protection (UMAP) method. Biological replicates were aggregated using canonical correlation analysis (CCA) method in Seurat. Dimension reduction, volcano plots, dot plots were generated using Seurat. All data is available on the gene expression omnibus (GEO) accession GEO: GSE269276.

**Supplemental Figure 1:**
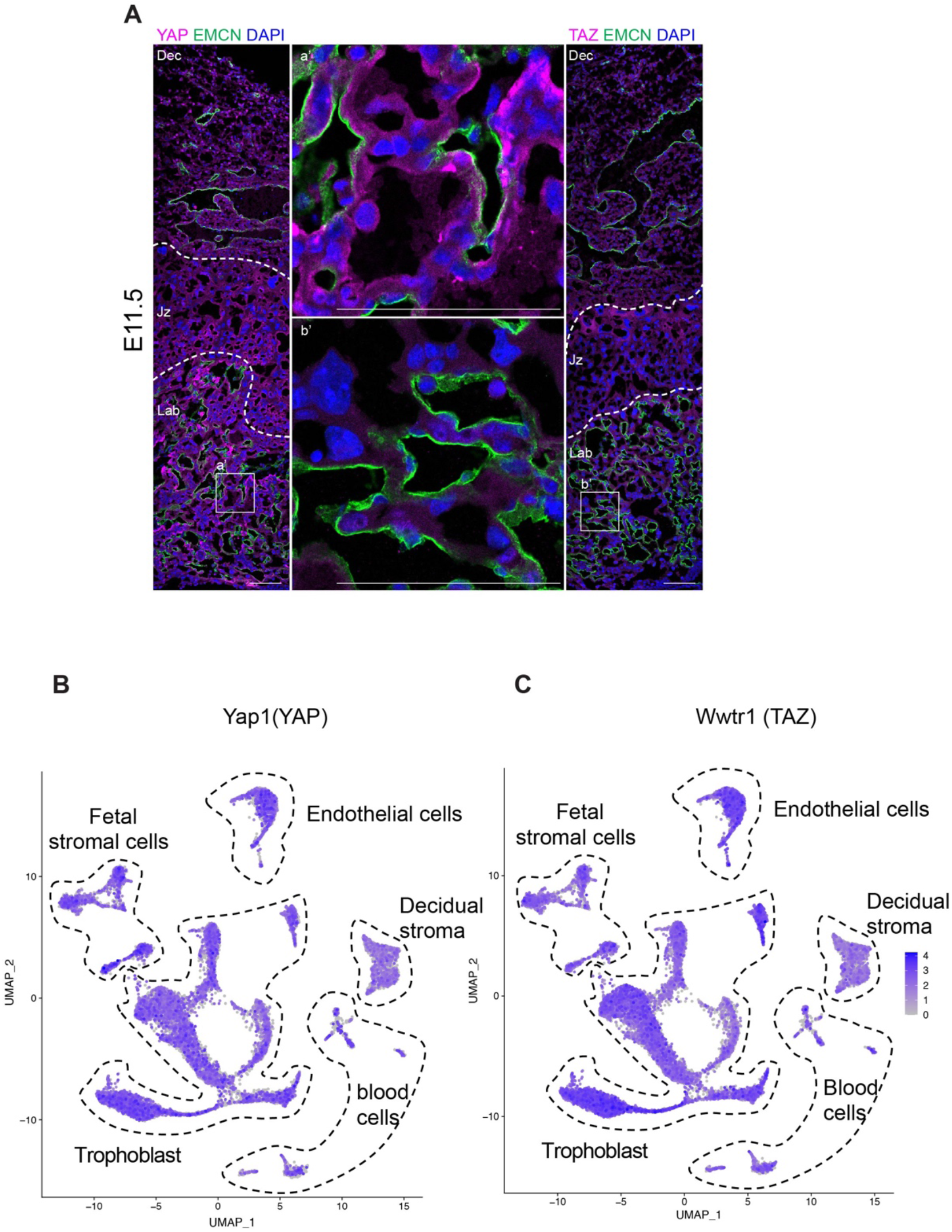
Expression of YAP and TAZ in the murine placental labyrinth. (A) Cross sections of E11.5 placenta containing the decidual (Dec), junction zone (Jz) and labyrinth (Lab) were immunostained for the endothelial cell marker Endomucin (green) and for YAP or TAZ (magenta) and were counterstained with the nucleic acid-binding dye DAPI (DAPI, blue). Images were generated by confocal microscopy. Boxed regions in the middle panels are shown magnified in the labyrinth region. Scale bars: 100 μm. (B,C) Feature plots of the expression of YAZ and TAZ from E9.5 to E14.5 placenta labyrinth single nuclei RNAseq dataset^27^.

**Supplemental Figure 2.**
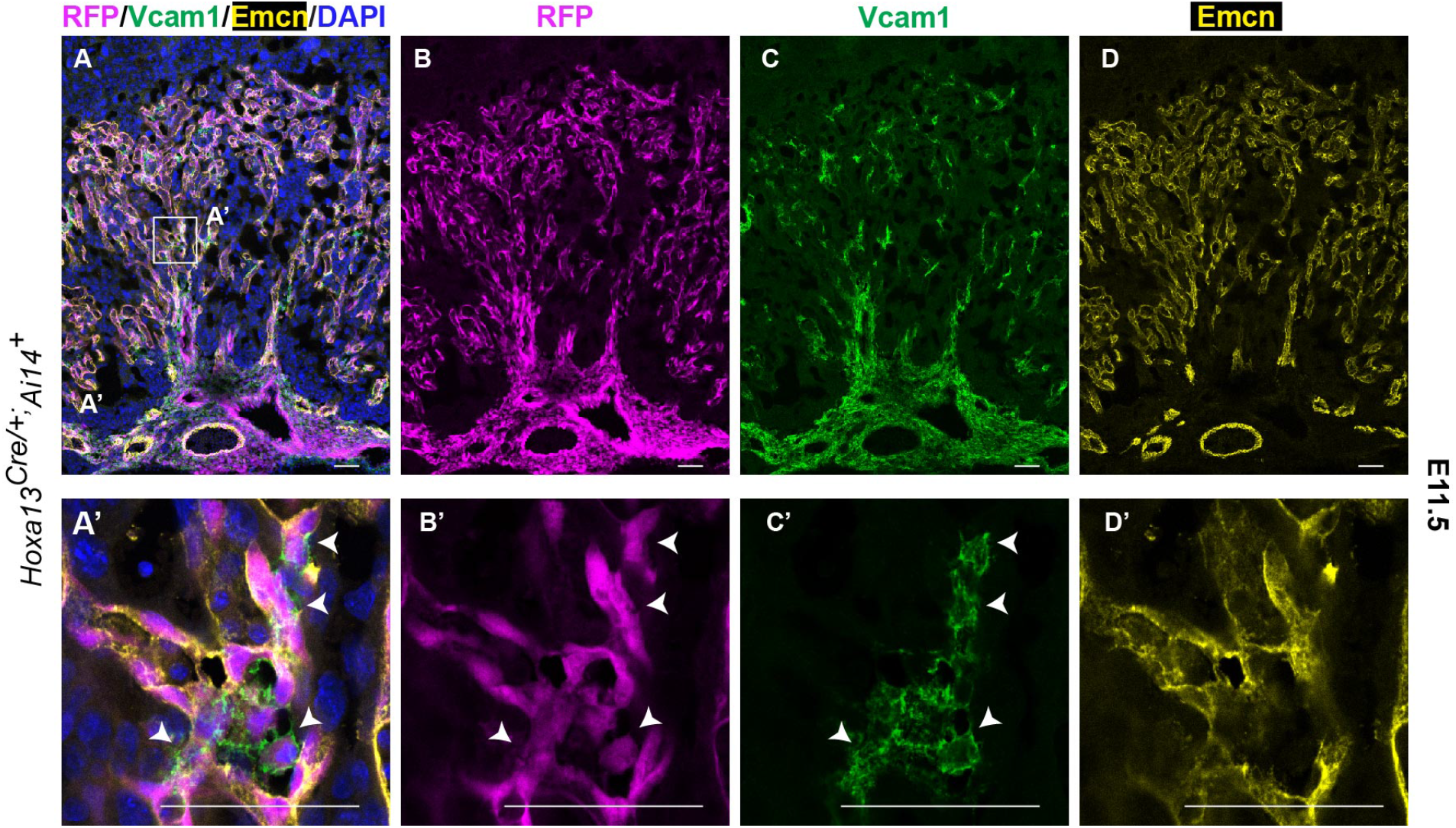
*Hoxa13*^*Cre*^ labels Emcn^+^ endothelial cells and VCAM1^+^ stromal cells in the placenta. (A-D) Representative images of E11.5 Hoxa13^Cre^; Ai14 placenta. (A-D) Immunofluorescence staining for RFP (magenta), VCAM1(green), Emcn (yellow), and DAPI (blue). (A’-D’) inserts of (A) the labyrinth region. Scale bars: 50μm.

**Supplemental Figure 3:**
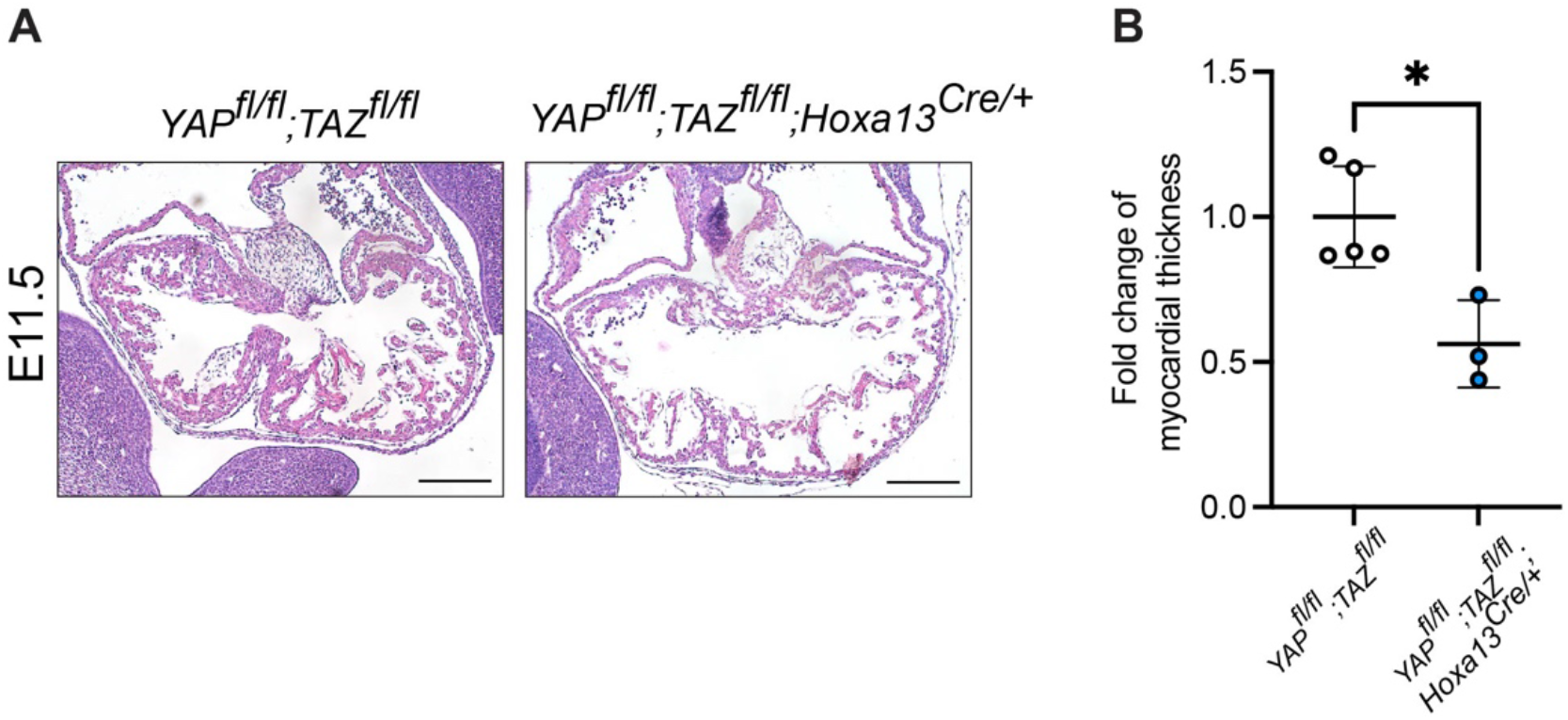
*YAP/TAZ*^*fl/fl*^; *Hoxa13*^*Cre/+*^ embryos exhibit thinned heart myocardium before death. (A). Littermate control *YAP/TAZ*^*fl/fl*^ and *YAP/TAZ*^*fl/fl*^; *Hoxa13*^*Cre/+*^ embryos were stained with H&E. n=3-4 per genotype. Scale bars: 200 μm. (B). Ratio of the difference in myocardial thickness of E11.5 littermate hears; 4 sections analyzed for each sample. Indicated *p<0.05 calculated by Student’s *t* test.

**Supplemental Figure 4:**
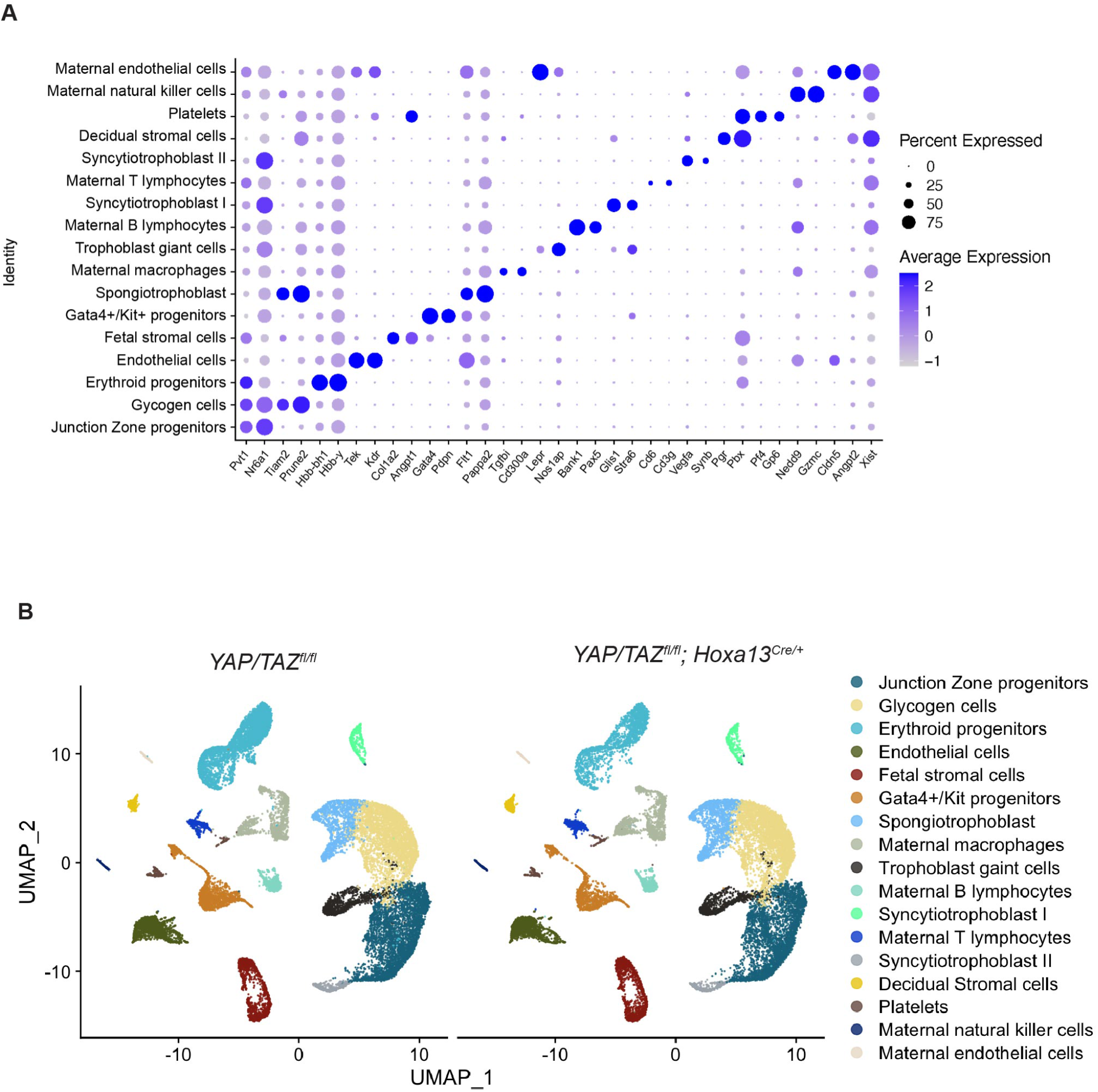
snRNAseq analysis reveals distinct cell populations in developing placental labyrinth. (A). Dot plot shows average expression and percent of nuclei in each cell cluster expressing canonical marker genes identified for each cluster. (B) UMAP dimension reduction analysis of snRNAseq data from *YAP*^*fl/fl*^*;TAZ*^*fl/fl*^ and *YAP*^*fl/fl*^*;TAZ*^*fl/fl*^; *Hoxa13*^*Cre/+*^ placentas.

**Supplemental Figure 5:**
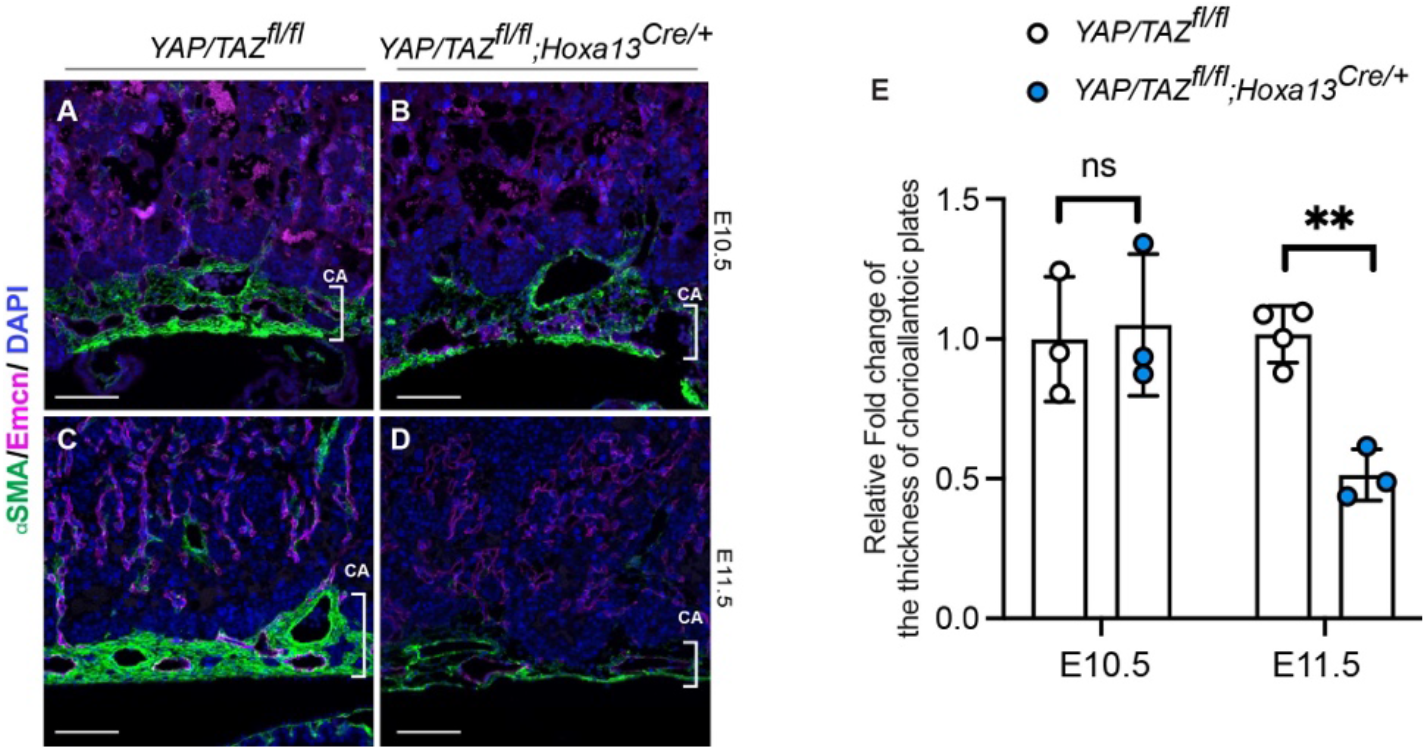
*YAP/TAZ*^*fl/fl*^; *Hoxa13*^*Cre/+*^ embryos exhibit impaired chorioallantoic plates. (A-D) Immunofluorescent images of E10.5 to E11.5 placenta were stained for endomucin (magenta) and for αSMA (green) and the nucleic acid-binding dye DAPI (blue). Scale bars: 200 μm. E. Quantification of the thickness of chorioallantoic plates; **p=0.011 calculated by Student’s *t* test.

